# Non-B-form DNA structures mark centromeres

**DOI:** 10.1101/209023

**Authors:** Sivakanthan Kasinathan, Steven Henikoff

## Abstract

Animal and plant centromeres are embedded in repetitive “satellite” DNA, but are thought to be epigenetically specified. To define genetic characteristics of centromeres, we surveyed satellite DNA from diverse eukaryotes and identified variation in <10-bp dyad symmetries predicted to adopt non-B-form conformations. Organisms lacking centromeric dyad symmetries had binding sites for sequence-specific DNA binding proteins with DNA bending activity. For example, human and mouse centromeres are depleted for dyad symmetries, but are enriched for non-B DNA and are associated with binding sites for the conserved DNA-binding protein CENP-B, which is required for artificial centromere function but is paradoxically non-essential. We also detected dyad symmetries and predicted non-B-form DNA structures at neocentromeres, which form at ectopic loci. We propose that centromeres form at non-B-form DNA because of dyad symmetries or are strengthened by sequence-specific DNA binding proteins. Our findings resolve the CENP-B paradox and provide a general basis for centromere specification.

## Introduction

Centromeres are nearly universally marked by the presence of nucleosomes containing the histone H3 variant CenH3 (CENP-A), which is sufficient to direct formation of the proteinaceous kinetochore that links chromosomes to the mitotic spindle (Mendiburo, et al. 2011). CENP-A is partitioned equally between sister chromatids during cell division and is deposited coincident with mitotic exit (Jansen, et al. 2007) by the conserved chaperone Holliday junction binding protein (HJURP/Scm3) (Kato, et al. 2007; Foltz, et al. 2009; Sanchez-Pulido, et al. 2009). Mechanisms dictating selection of particular loci as centromeres in complex genomes have remained elusive. One possibility is that centromeric DNA directs CenH3 deposition (De Rop, et al. 2012). Indeed, budding yeast ‘point’ centromeres are completely determined by ~120 bp sequences (Meraldi, et al. 2006; Gordon, et al. 2011). However, the centromeres of most other eukaryotes are made up of large tandem arrays of repetitive genetic elements whose role in centromere formation is unclear.

Centromeric repeats act as selfish genetic elements by driving non-Mendelian chromosome transmission during meiosis (Henikoff, et al. 2001) in turn spurring the rapid evolution of centromeric proteins to restore meiotic parity (Malik and Henikoff 2009). Incompatibilities between centromeric proteins and selfish DNA repeats may therefore serve as a molecular basis for speciation (Henikoff, et al. 2001). This centromere drive model is supported by observations confirming that centromeric alterations distort chromosome transmission in humans (Daniel 2002), mice (Chmatal, et al. 2014; Iwata-Otsubo, et al. 2017) and plants (Fishman and Saunders 2008). Critically, centromere drive supposes that centromeres are genetically defined at least through a transient stage of their evolution (Dawe and Henikoff 2006). However, the identification of neocentromeres that form at ectopic loci devoid of alphoid DNA (du Sart, et al. 1997) has led to the widely held view that centromere identity is specified independently of DNA sequence through “epigenetic” mechanisms involving CENP-A (Karpen and Allshire 1997; Ekwall 2007; Allshire and Karpen 2008).

Consistent with a role for genetic factors in centromere specification, primate centromeres have remained embedded in megabases of ~170-bp α-satellite tandem repeats over nearly 60 million years of evolution (Alexandrov, et al. 2001). The identification of CENP-B, a sequence-specific DNA-binding protein at centromeres that is highly conserved in mammals (Sullivan and Glass 1991), suggested a possible mechanism for DNA-encoded centromere specification (Masumoto, et al. 1989). However, although CENP-B is present in all primate genomes (Schueler, et al. 2010), the 17-nt CENP-B box sequence bound by the protein is not present on all centromeres within a species and is not found in all primates (Masumoto, et al. 1989; Haaf, et al. 1995). Further complicating this “CENP-B paradox” (Goldberg, et al. 1996; Kipling and Warburton 1997), CENP-B binding is required for *de novo* centromere formation on artificial chromosomes (Ohzeki, et al. 2002) and has been shown to enhance the fidelity of chromosome segregation (Fachinetti, et al. 2015). Reconciling these observations concerning variation in centromeric DNA with evidence that centromeres are epigenetically defined remains an outstanding challenge.

Here, we reconsider the role of DNA sequence in specification of centromere identity. We mined publicly available whole-genome sequencing datasets for centromeric DNAs from great apes, Old World Monkeys (OWMs), mouse, chicken, plants, and yeasts and characterized clade-specific variation in abundance of dyad symmetries predicted to adopt non-B-form DNA conformations. We found a highly restricted distribution of CENP-B boxes limited to great apes and mouse. Indeed, we could show that absence of CENP-B boxes in OWM α-satellite corresponds to lack of CENP-B binding. We discovered that this loss of CENP-B binding was correlated with an increased tendency of centromeric satellites to form predicted non-B-form DNA structures such as cruciforms. Using experimental datasets, we were able to detect these non-B-form DNA structures at functional human and mouse centromeres. In budding yeasts, we identified a similar association between centromeric dyad symmetry and exaptation of a CENP-B -like DNA-binding protein. We also predicted non-B-form DNA formation at the human Y chromosome centromere and human and chicken neocentromeres, which are not associated with CENP-B binding. Based on these data, we advance a unifying model for centromere specification based on recognition of non-B-form DNA structures, either aided or unaided by sequence-specific DNA binding proteins.

## Results

### Dyad symmetries are common features of centromeres

Sequence analysis of centromeres has been challenging because of the intractability of repeats; however studies in the fission yeast *Schizsaccharomyces pombe*, which has regional centromeres similar to vertebrates, and analyses of primate α-satellite consensus sequences and budding yeast centromeres have suggested the existence of sequence-encoded features that are important for centromere function (Koch 2000; Catania, et al. 2015). To identify potential DNA sequence determinants that might have been overlooked in recent studies of centromere specification, we first catalogued <10-nt dyad symmetries. Using published consensus sequences or libraries of species-specific satellite consensus sequences derived from *de novo* tandem repeat detection (**Figure S1A**), we scanned deep (>10X coverage) publicly available paired-end, whole genome sequencing datasets from a sampling of vertebrates, fission yeast, and plants to identify centromeric sequences, which are typically poorly represented in genome assemblies. In chicken, which has both repetitive and unique centromeres (Shang, et al. 2010), we restricted our analyses to well characterized unique sequences. Visual examination of centromeric sequences identified dyad symmetries with species-specific variation (**Figure 1A**). For species with high-quality genome assemblies, we compared dyad symmetries in centromeric regions and dinucleotide composition-matched background genomic regions without known centromeres and confirmed the pattern of species-specific enrichment of dyad symmetries (**Figure 1B**). To determine whether dyad symmetries may have been selected for during centromere evolution, we compared centromeric sequences from each species to random permutations of the same sequences to account for nucleotide composition. Enumeration of dyad symmetries over varying palindrome lengths revealed enrichment of >3-bp dyad symmetries in OWMs, horse, chicken, stickleback, fission yeast, and plants, but not in great apes or mouse (Figure 1C, **S2A**). Based on these analyses, we conclude that dyad symmetries are a unique feature of many eukaryotic centromeres and may have been selected for during centromere evolution.

**Figure 1.**
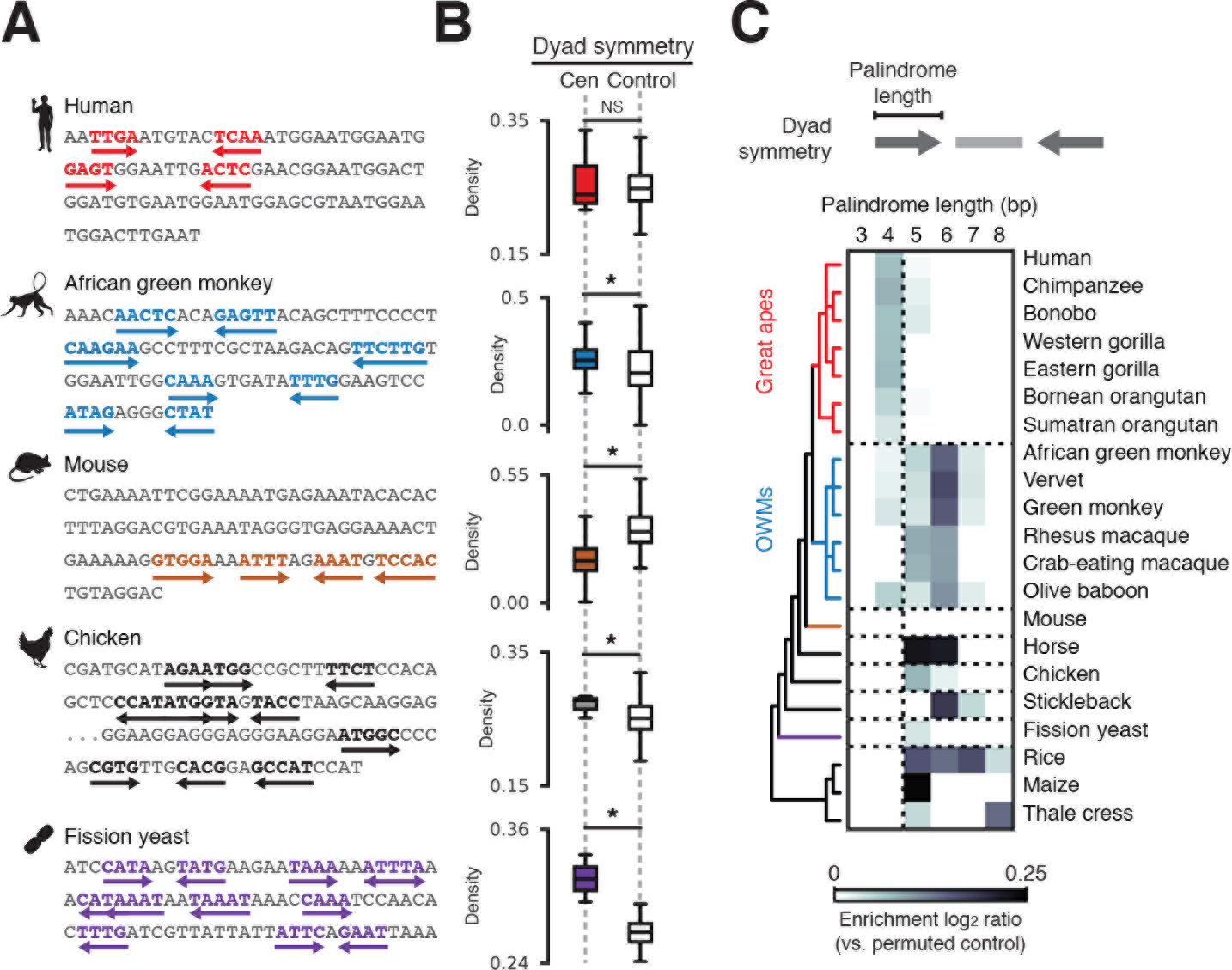
Patterns of DNA dyad symmetry at eukaryotic centromeres. **(A)** Examples of dyad symmetries in centromeric DNA sampled randomly from human, African green monkey, mouse, chicken, and fission yeast whole-genome sequencing datasets. **(B)** Dyad density, which is defined for a given sequence as the total number of palindromic positions with palindrome length > 4 and spacer length < 20 normalized by sequence length, at centromeres relative to composition-matched background genomic regions. Asterisks indicate two-sample Kolmogorov-Smirnov *p* < 0.05. **(C)** Enrichment (relative to permuted sequence) of dyad symmetries over varying palindrome lengths in read ends mapping to centromeres or from sequences sampled from genome assemblies for a variety of organisms. The displayed phylogeny is based on NCBI Taxonomy annotations.

### Predicted non-B-form DNA structures at dyad-enriched centromeres

Given the possibility that regions of dyad symmetry may adopt non-B DNA conformations (Pearson, et al. 1996) and reports of stem-loop, hairpin, and triplex nucleic acid at human centromeres (Ohno, et al. 2002; Jonstrup, et al. 2008; Garavis, et al. 2015; Aze, et al. 2016), we sought to characterize the theoretical secondary structure formation potential of centromeric DNAs. We used a computational method that models stress-induced structural transitions (SIST) in DNA (Zhabinskaya, et al. 2015) to determine whether variation in dyad symmetry corresponds to differing predispositions for adopting non-B conformations. Comparing SIST DNA melting and cruciform extrusion scores for centromeric sequences and dinucleotide composition-matched non-centromeric background genomic intervals for select species, we found that dyad-enriched centromeres were associated with significantly higher levels of predicted non-B-form DNA (**Figure 2A**). We then used an independent approach to predict folding free energies for centromeric DNAs and found that species predicted by SIST to adopt non-B-form DNA tended to form more stable secondary structures (**Figure 2B**). For example, consistent with the SIST predictions, the distributions of predicted free energies for dyad-enriched α-satellite from OWMs were substantially left-shifted compared to dyad-depleted great ape α-satellite, suggesting that OWM centromeres may adopt more stable secondary structures than great ape centromeres (two-sample Kolmogorov-Smirnov *p*-value ≪ 1e-5; **Figures 2B,C**, **S2B**). Similar trends were observed in other species (**Figure 2C**). From these analyses, we conclude that centromeres enriched for dyad symmetries may adopt stable non-B-form DNA structures such as cruciforms.

**Figure 2.**
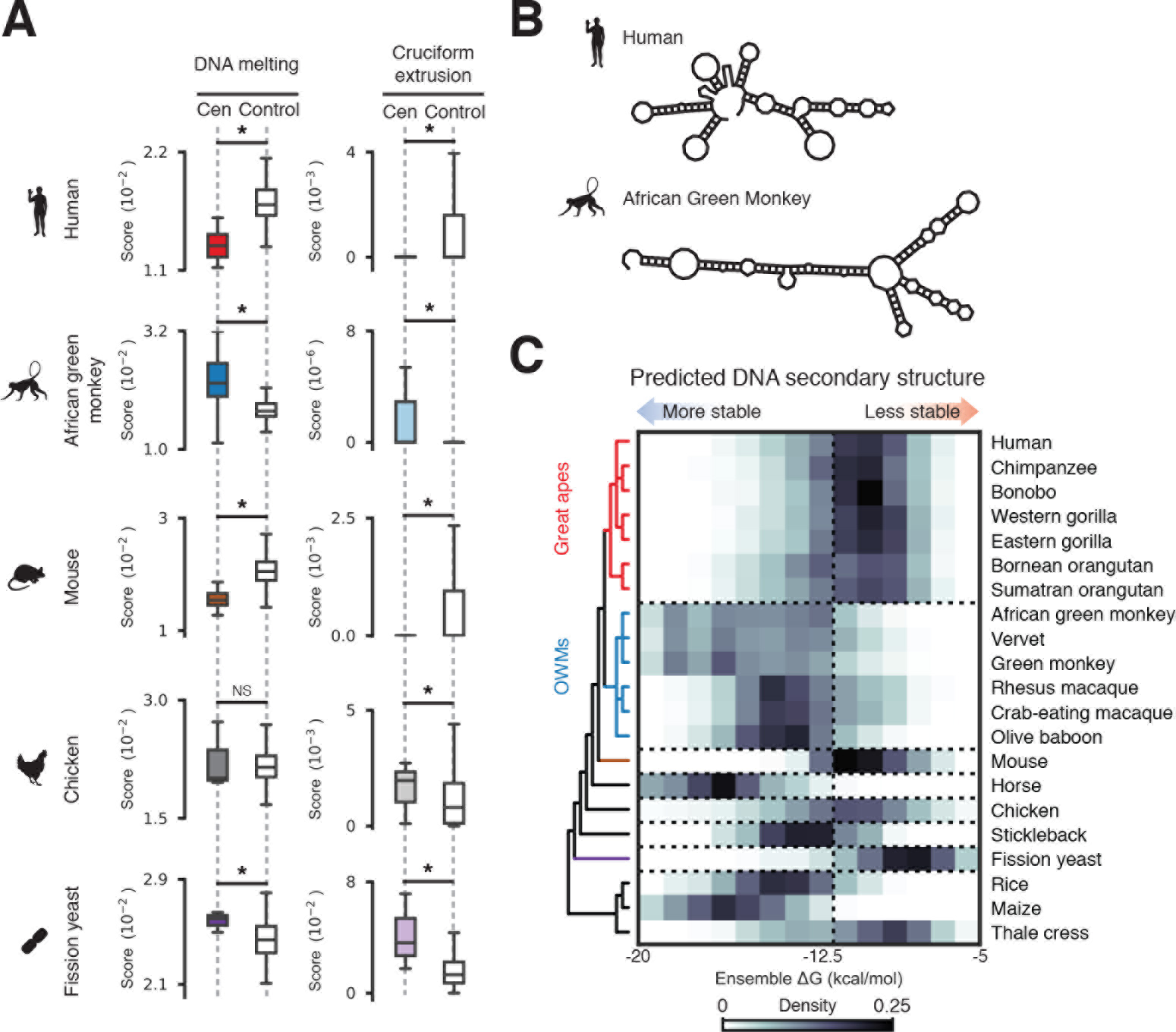
Centromeric dyad symmetries are predicted to adopt non-B-form structures. **(A)** Scores from stress-induced structural transition (SIST) model predictions of DNA melting (left) and cruciform extrusion (right) for centromeric sequences and composition-matched background genomic regions from human, African green monkey, mouse, chicken, and fission yeast genomes. Asterisks indicate two-sample sample Kolmogorov-Smirnov *p* < 0.05. (*B*) Examples of minimum free energy secondary structure predictions for randomly selected α-satellite monomers from human and African green monkey. (**C**) DNA secondary structure folding free energy predictions for read ends mapping to centromeres or from sequences sampled from genome assemblies from the indicated species. The displayed phylogeny is based on NCBI Taxonomy annotations.

### CENP-B-associated enrichment of non-B-form DNA at dyad-depleted human and mouse centromeres

We next sought to verify computational structure predictions using publicly available data from genome-wide mapping of non-B-form DNA using potassium permanganate treatment (permanganate-seq) in mouse and human cells (Kouzine, et al. 2013; Kouzine, et al. 2017). In mouse, minor satellite (MiSat) sequences constitute the functional centromere (Joseph, et al. 1989), while adjacent major satellite (MaSat) domains are heterochromatic (Horz and Altenburger 1981). We confirmed the relative abundances of these satellites (**Figure 3A**) and, in agreement a previous report (Guenatri, et al. 2004), we detected CENP-B boxes in MiSat arrays while MaSat sequences contained very few CENP-B boxes (**Figure 3A**). Importantly, unlike centromeric MiSat sequences, heterochromatic MaSat arrays were predicted to more favorably adopt non-B-form structures (**Figure 3B**). MaSat sequences were enriched for permanganate-seq reads concordant with structure predictions (**Figure 3C**). Surprisingly, MiSat sequences were also highly enriched for permanganate-seq signal (**Figure 3C**). Permanganate-seq signal increased in activated B-cells, which are undergoing cell division, relative to quiescent resting B-cells (**Figure 3C**). To determine whether non-B-form DNA was associated with functional MiSat sequences occupied by CENP-A, we analyzed chromatin immunoprecipitation and sequencing (ChIP-seq) data (Iwata-Otsubo, et al. 2017). CENP-A occupancy was positively correlated with permanganate-seq signal (Spearman’s ρ = 0.45; *p* ≪ 1e-10), with MiSat sequences associated with low-scoring CENP-B sites having the least CENP-A and permanganate-seq alignments (**Figure 3D,E**).

**Figure 3.**
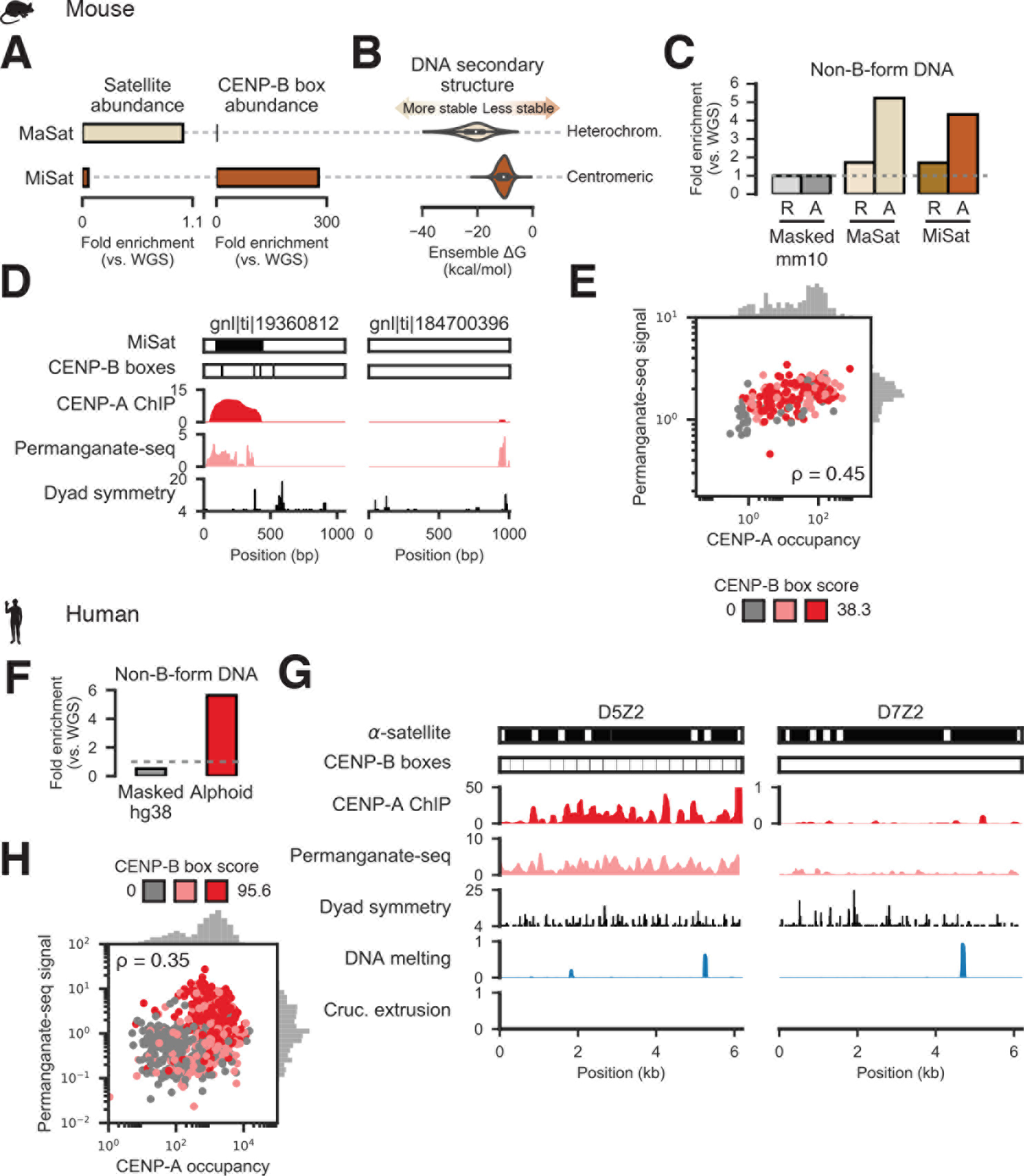
Non-B-form DNA detected experimentally at dyad-depleted functional human and mouse centromeres. (**A**) Abundance of heterochromatic major (MaSat) and centromeric minor (MiSat) satellite 22 fragments (top) and the minimal CENP-B box (middle) in mouse whole-genome sequencing reads. (**B**) DNA secondary structure free energy distributions from computational prediction for MaSat and MiSat-containing Sanger reads. (**C**) Fraction of permanganate-seq reads from resting (R) and LPS-activated (A) cells mapping to the masked mouse genome (Masked mm10) or Sanger reads containing MiSat or MaSat monomers. (**D**) Examples of Sanger reads harboring MiSat and CENP-B box sequences (left) or devoid of predicted centromeric features (right) and signal from CENP-A ChIP-seq, permanganate-seq, and dyad symmetry analysis. (**E**) Correlation between total permanganate-seq signal from activated B cells (normalized to control) and input-normalized CENP-A occupancy for MiSat-containing Sanger reads. Scatter plot points are colored based on CENP-B box score tertile, where scores are defined as the sum of scores for all FIMO-defined CENP-B boxes occurring on a read. (**F**) Fraction of permanganate-seq reads aligning to the repeat-masked hg38 assembly and HuRef Sanger alphoid reads normalized to number of reads from whole-genome sequencing of HuRef mapping to the respective assemblies. (**G**) Examples of permanganate-seq, CENP-A ChIP, dyad symmetry, and predicted DNA melting and cruciform transition probabilities for functionally active (D5Z2) and inactive (D7Z2) α-satellite repeat arrays. Note that the CENP-A ChIP tracks are on different scales. (**H**) Correlation between total permanganate-seq signal and CENP-A occupancy for alphoid Sanger reads. Scatter plot points are colored based on CENP-B box score tertile.

Similarly, contrary to the predicted poor tendency for human alphoid DNA to adopt non B-form DNA structures (**Figure 2C**), more than half the human permanganate-seq data aligned to α-satellite, representing a ~10-fold enrichment for alphoid DNA (**Figure 3F**). Regions enriched for non-B-form DNA appeared to colocalize with CENP-A, particularly at functional α-satellite dimers with CENP-B boxes that we previously identified by ChIP-seq (Henikoff, et al. 2015), but were depleted at presumably non-functional α-satellite arrays that are not at centromeres (Slee, et al. 2012) and not associated with CENP-B boxes (**Figure 3G**). Indeed, CENP-A occupancy and permanganate-seq signal (Spearman’s ρ = 0.35; *p* ≪ 1e-10), with alphoid arrays associated with low-scoring CENP-B sites having the least CENP-A and permanganate-seq signal (**Figure 3H**). We conclude that non-B-form DNA is characteristic of functional, CENP-B-associated human and mouse centromeric satellites and may form in a cell cycle-dependent manner.

### Functional Old World Monkey centromeres enriched for non-B-DNA are not bound by CENP-B

Given the inverse relationship between CENP-B binding and non-B DNA, we speculated that CENP-B boxes might be specific to great-ape centromeres. To test this hypothesis, we searched for matches to the multiple alignment-based consensus CENP-B box sequence (TTCGNNNNANNCGGG) required to support CENP-B DNA binding (Iwahara, et al. 1998) in randomly sampled whole-genome sequencing reads. Although OWMs tended to have more α-satellite (**Figure 4A**), CENP-B boxes were highly enriched in great apes whereas OWM genomes contained negligible matches to the minimal CENP-B box (**Figure 4B**), consistent with a previous report (Goldberg, et al. 1996). We further verified great ape-specific enrichment of CENP-B boxes using the SELEX-defined CENP-B motif (Jolma, et al. 2013) (**Figure S1B**). To more finely characterize functional centromeric sequences in divergent primates, we mapped genomic binding of CENP-A and CENP-B in K562 (human) and Cos-7 (African green monkey) cell lines using CUT&RUN, which profiles DNA fragments released by antibody-targeted nuclease cleavage *in situ* (Skene and Henikoff 2017b). We detected substantial enrichment for alphoid sequence in CENP-A CUT&RUN in both human and African green monkey (**Figure 4B**). Consistent with a paucity of CENP-B boxes in OWMs, CENP-B protein binding within alphoid sequence was observed in human but not African green monkey (**Figure 4C**). CENP-B boxes were highly enriched in CENP-A CUT&RUN reads from K562 cells, but depleted in CENP-A associated sequences from Cos-7 cells (**Figure 4D**). Taken together, these analyses suggest that CENP-B protein and binding sites are specific to dyad-depleted great ape centromeres.

**Figure 4.**
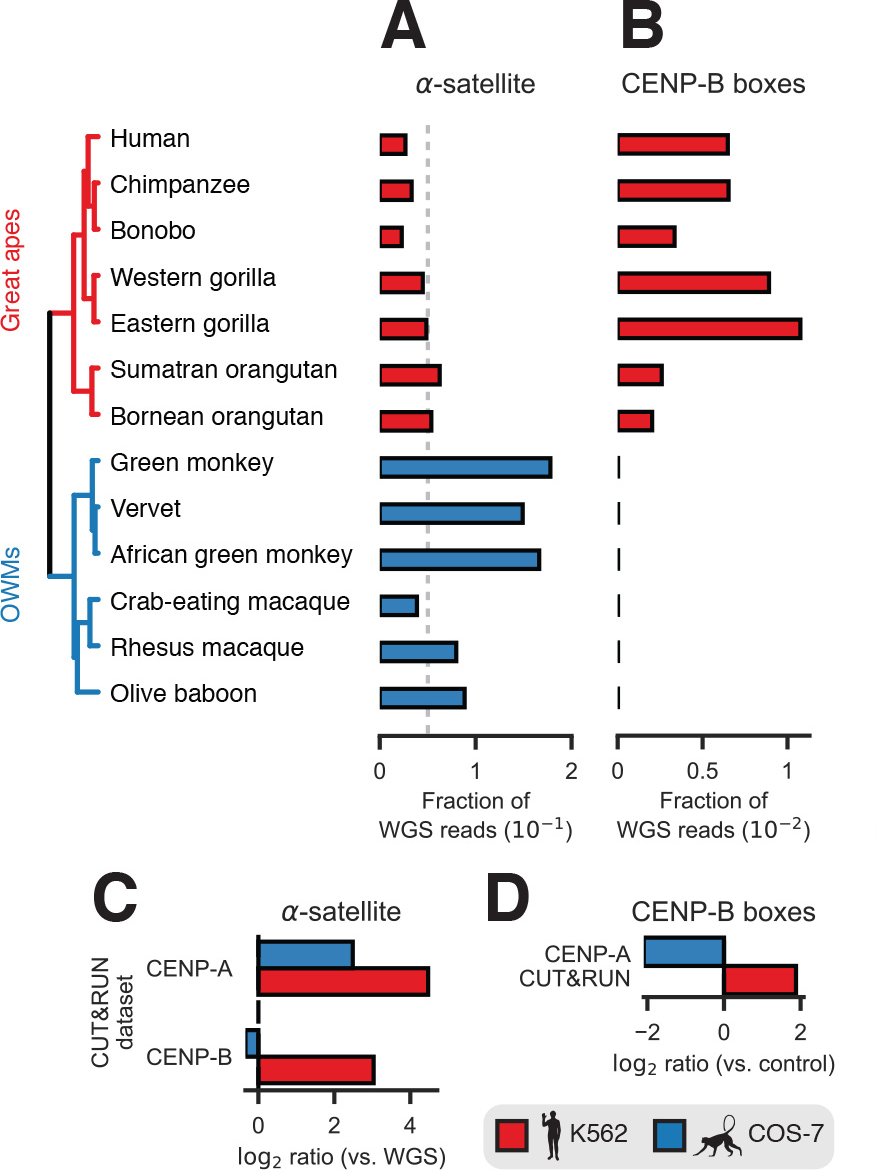
Primate CENP-B binding is restricted to dyad-depleted great ape centromeres. **(A)** Estimated abundance of α-satellite sequences in a sampling of simian primates. (**B**) Enrichment of minimal CENP-B box sequences in raw reads from whole-genome sequencing. The displayed phylogeny is a chronogram based on mitochondrial genomes and is adapted from the 10kTrees Project (Arnold, et al. 2010). Abundance of α-satellite sequence in aligned reads (**C**) and abundance of matches to the minimal CENP-B box sequence in raw reads (**D**) in CUT&RUN experiments performed in human (K562) and African green monkey (Cos-7) cell lines relative normalized to α-satellite abundance in whole-genome sequencing reads.

### CENP-B-depleted human Y centromere and vertebrate neocentromeres are enriched for non-B-form DNA

The observations that the human Y chromosome centromere and neocentromeres are devoid of CENP-B boxes (Haaf, et al. 1995; Saffery, et al. 2000) and that the CENP-B protein is non-essential (Kapoor, et al. 1998) contradict the requirement for CENP-B in *de novo* centromere assembly (Ohzeki, et al. 2002). To reexamine this “CENP-B paradox” (Goldberg, et al. 1996; Kipling and Warburton 1997) in light of DNA-encoded structural features at centromeres, we asked whether DYZ3 alphoid repeats from the human Y chromosome and vertebrate neocentromere sequences are associated with non-B DNA. First, we compared predicted folding free energies of fragments derived from DYZ3 and non-DYZ3 alphoid DNA and found that DYZ3 sequences are associated with thermodynamically favorable non-B structures (**Figure 5A,B**), suggesting that the CENP-B binding may not be required for *de novo* centromerization. Next, we analyzed CENP-A ChIP-seq data from human cell lines bearing neocentromeres (Hasson, et al. 2013) and detected enrichment for dyad symmetries predicted to form stable secondary structures in CENP-A-associated neocentromere domains (**Figures 5C**). To determine whether cruciform structures, a specific class of non-B-form DNA, are associated with neocentromeres, we used data from genome-wide analysis of palindrome formation (GAP-seq) based on DNA renaturation and S1-nuclease treatment (Yang, et al. 2014), and found regions that may form cruciforms *in vivo* at neocentromeres (**Figures 5C**). Neocentromeres were also markedly enriched for dyad symmetries relative to base composition-matched randomly selected genomic regions and native centromeric sequences (**Figure 5C**). We also analyzed data from a chicken neocentromere cell line to determine whether a similar trend is generally observed in vertebrates. Like other mammalian centromeres, chicken neocentromeres were also enriched for short dyad sequences and predicted to undergo strand separation and cruciform transitions (**Figure 5D**). Taken together, these analyses suggest that native centromeres depleted for CENP-B boxes such as the human Y chromosome centromere and neocentromeres are enriched for non-B-form DNA structures.

**Figure 5.**
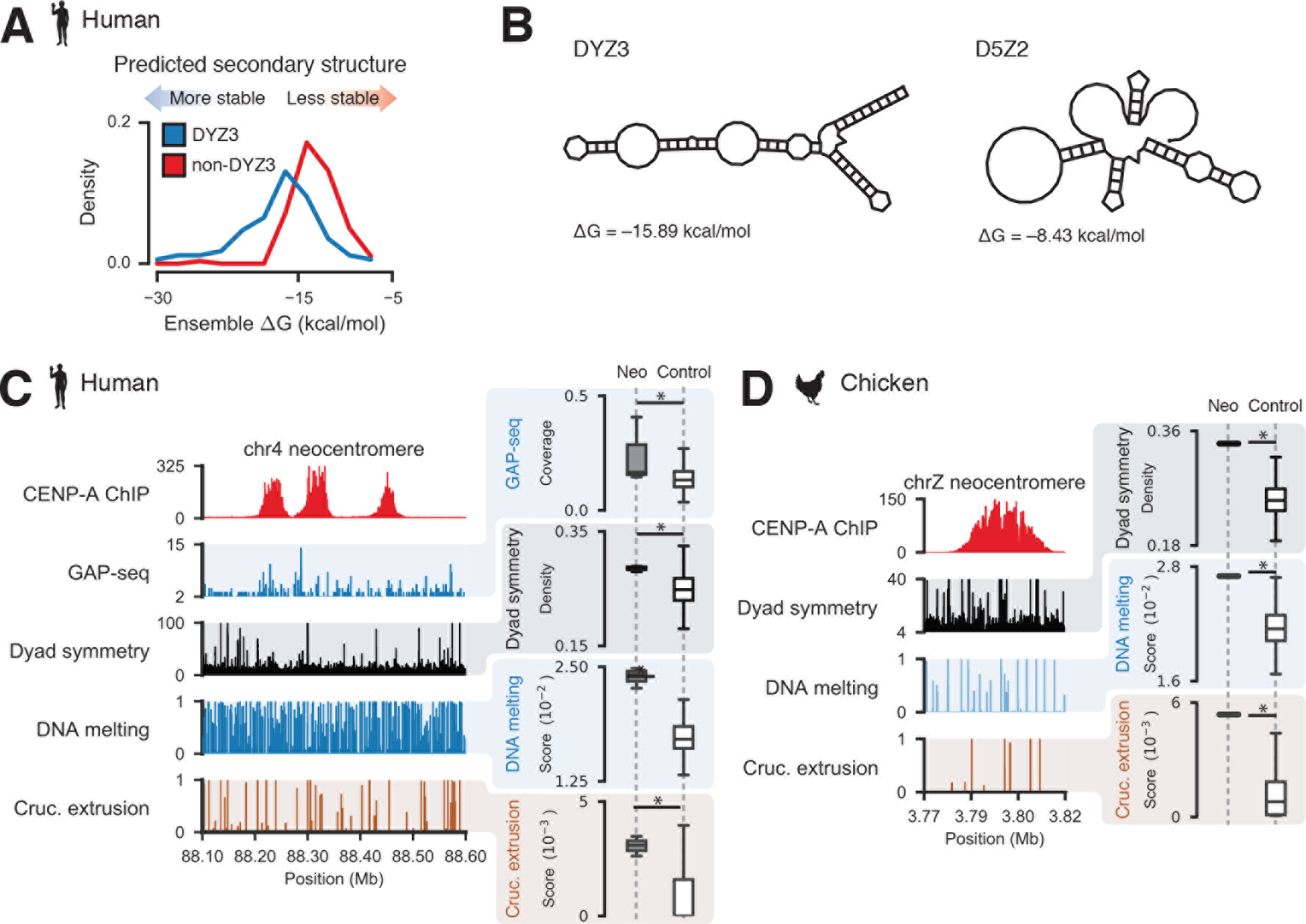
The human Y centromere and vertebrate neocentromeres are associated with dyad symmetries and non-B-form DNA. (**A**) Predicted ensemble free energies for DYZ3 and non-DYZ3 alphoid satellites. (**B**) Examples of minimum free energy structures for a DYZ3 and D5Z2 alphoid fragments. A human chromosome 13 neocentromere (**C**) and a chicken chrZ neocentromere (**D**) with profiles from CENP-A ChIP-seq and SIST-predicted DNA melting and cruciform extrusion probabilities (left panels). Dyad symmetry and SIST DNA melting and cruciform extrusion scores for neocentromeres (“neo”) and composition-matched non-centromeric background genomic intervals (right panels). Data 23 from genome-wide analysis of palindrome formation with sequencing (GAP-seq), which was performed in human cell lines, is also included in (**A**). Asterisks indicate two-sample Kolmogorov-Smirnov *p* < 0.05.

### Exaptation of a CENP-B-like protein at dyad-depleted budding yeast centromeres

In contrast to most eukaryotes, the saccharomycetes have among the simplest known centromeres, which are fully determined by a ~120 bp sequence composed of three centromere determining elements (CDEs I-III) (Clarke and Carbon 1985). To gain insight into whether dyad symmetries are a conserved feature of centromeres in diverse eukaryotes, we analyzed sequences of well-characterized budding yeast centromeres. We determined the extent of CENP-A binding at annotated centromeres using published datasets (Henikoff, et al. 2014; Thakur, et al. 2015) and quantified the extent of DNA melting and cruciform transition predicted by SIST (Zhabinskaya, et al. 2015). We first analyzed the average ChIP-seq signal for the CENP-A homologue Cse4 (Henikoff, et al. 2014), dyad symmetry, and SIST DNA melting and cruciform extrusion scores from *Saccharomyces cerevisiae*. Similar to what we observed in vertebrates, we found higher levels of predicted non-B DNA at *S. cerevisiae* centromeres that were enriched for dyad symmetries compared to composition-matched non-centromeric genomic regions (**Figures 6A and S6B**). We detected a similar pattern in enrichment for dyad symmetries and non-B-form DNA at the centromeres of other *sensu strictu* yeasts; however, despite similar sequence composition to *sensu strictu* saccharomycetes the *sensu lato* species *S. castellii* and *S. darienensis* had comparatively less dyad symmetry and lower SIST DNA melting and cruciform extrusion scores (Figures 6B,C **and S3C,D**). Recently, *S. castellii* and *S. dairenensis* were shown to have divergent point centromeres (Kobayashi, et al. 2015), with a substantially different CDEI region devoid of a binding site for the basic helix-loop-helix transcription factor Cbf1, which is found at CDEI sequences of *sensu strictu* centromeres (**Figure 6B,C**). We found that the consensus site at CDEI of dyad-depleted *sensu lato* yeasts is strongly predicted (*p* < 1e-5) to bind the DNA-bending general regulatory factor Reb1 (Figures 6B,C **and S3E**) based on searching a database of 203 yeast transcription factors (MacIsaac, et al. 2006) and comparison to a consensus motif from high-resolution mapping (Kasinathan, et al. 2014). These analyses suggest that non-B-form DNA at centromeres may represent an ancient mechanism for centromere specification in eukaryotes and that DNA-binding proteins such as CENP-B and Reb1 may serve an important role in shaping the evolution of centromeric DNA.

**Figure 6.**
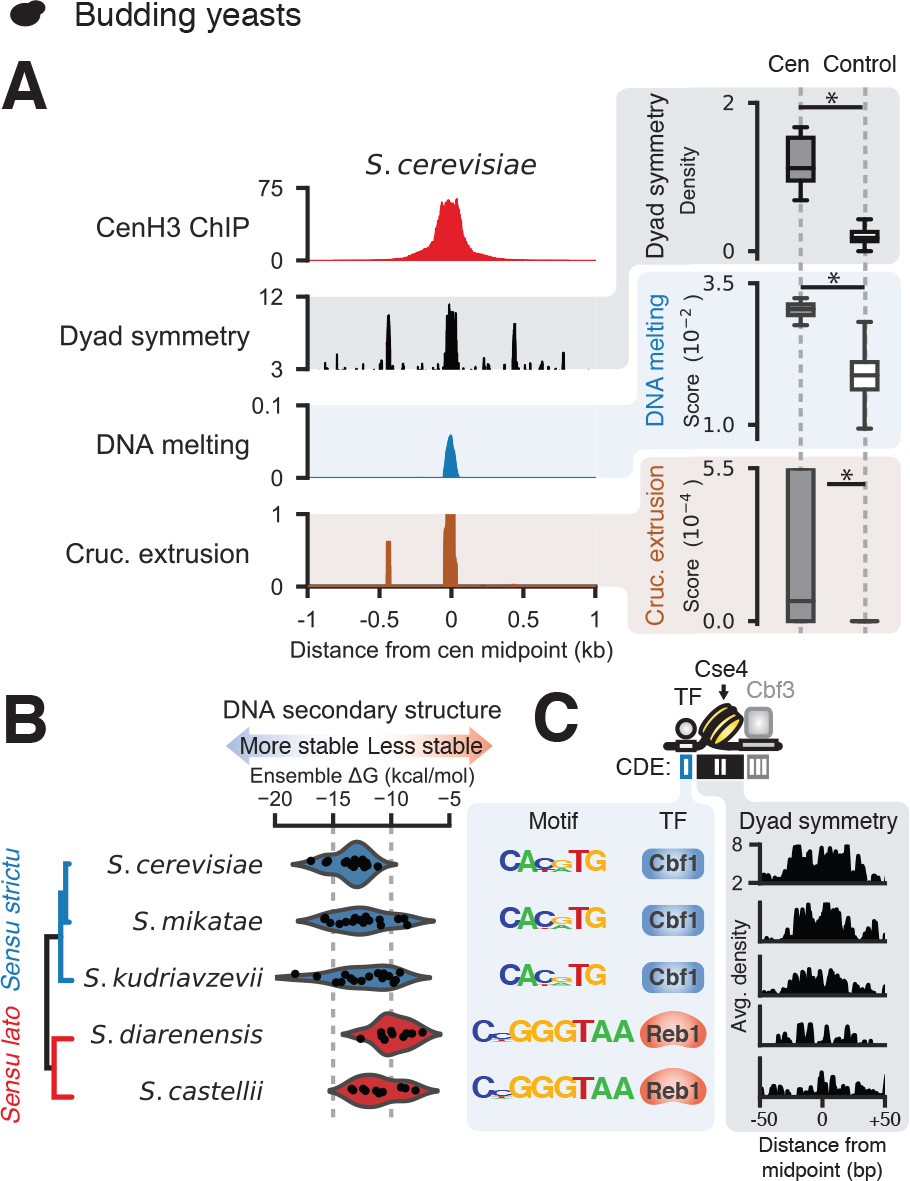
Centromeric dyad symmetries are features of yeast centromeres. Average CenH3 signal, dyad symmetry, and SIST melt and cruciform profiles (left panels) and comparison of dyad densities and SIST melt and cruciform scores for centromeres versus nucleotide composition-matched, randomly selected genomic intervals (right panels) in *S. cerevisiae* (**A**). (**B**) Predicted ensemble free energy distributions for centromeric sequences from *sensu strictu* and *sensu lato* saccharomycetes with well-annotated genomes. (**C**) Enriched CDEI motifs for saccharomycetes and average estimated dyad densities over CDEII.

## Discussion

We used comparative analysis of whole genome sequencing and functional genomic datasets to define evolutionary transitions in centromeres (**Figure 7A**). We found that short dyad symmetries that are predicted to adopt non-B-form structures are characteristic of OWM, chicken, and fission yeast centromeres, while great ape and mouse centromeres were comparatively depleted for dyad symmetries and not predicted to form non-B DNA. Surprisingly, both human and mouse centromeres were found to be associated with non-B DNA *in vivo*, with greater enrichment for non-B-form DNA at CENP-A-occupied satellite sequences associated with CENP-B boxes. Importantly, we did not detect CENP-B boxes at the OWM centromeres, which have α-satellite repeats predicted to adopt stable non-B-form structures. We also profiled CENP-A and CENP-B binding in human and African green monkey cell lines and demonstrated directly that functional OWM centromeres are not bound by CENP-B. Further, we found that the human Y chromosome centromere, which notably does not bind CENP-B, is predicted to form more thermodynamically favorable non-B-form DNA structures than other human centromeres. These observations resolve conflicting reports about the presence of CENP-B boxes in OWMs (Goldberg, et al. 1996; Yoda, et al. 1996).

**Figure 7.**
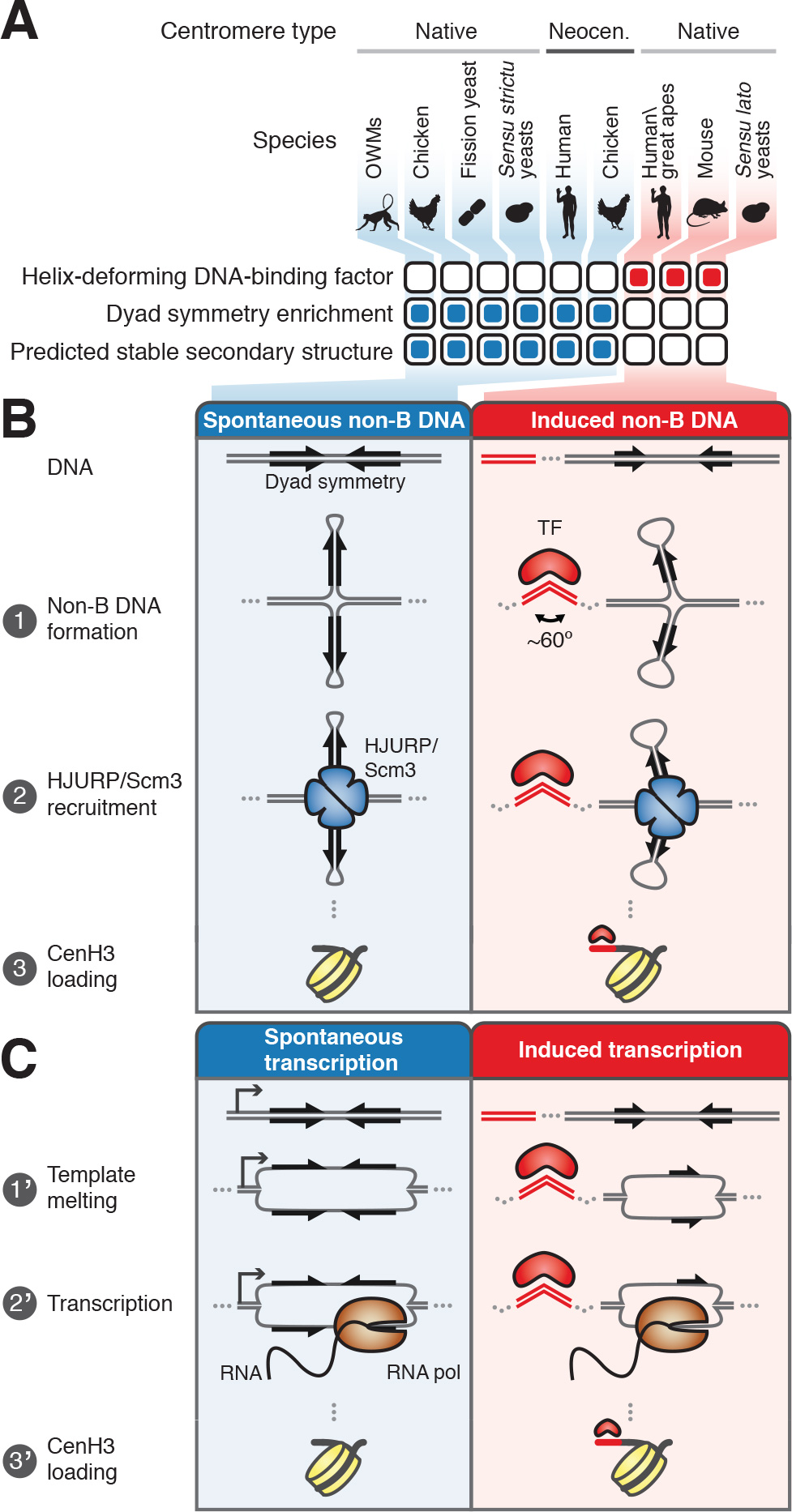
Models for genetic centromere specification. (**A**) Summary of centromeric DNA sequence type, association with helix-deforming DNA binding protein, dyad symmetry, and predicted secondary structure forming tendency for various eukaryotes. (**B**) Repetitive centromeres vary in their predilection for forming cruciform structures exemplified by alphoid sequences of Old World Monkeys, which are predicted to form stable non-B DNA structures, and great apes, which do not preferentially adopt non-B DNA structures. In great apes, CENP-B binding may facilitate formation of non-B-form DNA such as cruciforms. Cruciform structures are recognized by HJURP/Scm3 chaperones, which deposit CENP-A nucleosomes. (**C**) Alternatively, OWM AS units may be spontaneously transcribed, while CENP-B binding may facilitate transcription of great ape alphoid units, with the RNAs contributing to deposition of CENP-A.

Consistent with the presence of non-B-form DNA structures at functional centromeres, single-stranded DNA, hairpins, triplexes, and i-motifs have been observed in α-satellite *in vitro* and/or *in vivo* (Ohno, et al. 2002; Jonstrup, et al. 2008; Garavis, et al. 2015; Aze, et al. 2016). Taken together with our genomic analyses, these lines of evidence suggest testable models for the specification of centromere identity (**Figure 7B,C**). One possibility is that non-B-form DNA directly specifies centromere identity (**Figure 7B**). Because the CENP-A chaperone HJURP was named based on its *in vitro* Holliday junction-binding activity (Kato, et al. 2007), it is tempting to speculate that the four-way junction DNA structures recognized by HJURP are short cruciforms, which may form spontaneously or inducibly. Organisms such as OWMs may have satellites capable of adopting energetically favorable conformations recognized by HJURP and its ortholog Scm3, which is the CenH3 chaperone in both budding and fission yeast. In contrast, other species such as great apes may require the binding of a sequence-specific DNA-binding protein such as CENP-B to promote formation of non-B DNA structures. It is intriguing that both mammalian CENP-B and yeast Reb1 bend DNA ~60°, raising the possibility that formation of non-B-form DNA in activated mouse and human B cells is directly or indirectly mediated by CENP-B-mediated DNA bending.

Alternatively, centromere specification may occur through a transcription-based mechanism that produces RNAs with secondary structure (**Figure 7C**). Transcription may occur readily at centromeres that adopt non-B-form structures, which include melted DNA, while sequences that are comparatively resistant to adopting non-B conformations may require the action of an architectural DNA-binding protein such as CENP-B or Reb1. Transcription in the case of dyad-enriched satellites may also occur through recognition by the rDNA transcription factor UBF, which has cruciform-binding activity (Copenhaver, et al. 1994), and RNA polymerase (Pol) I, whereas transcription at sequences that do not favorably adopt non-B structures could occur via RNA Pol II. In support of this model, Pol II transcription has been shown to occur at human centromeres (Quenet and Dalal 2014; McNulty and Sullivan 2017) and to be functionally important in budding yeast centromeres (Ohkuni and Kitagawa 2011). Pol I has similarly been suggested to be involved in human centromere function (Wong, et al. 2007).

These non-mutually exclusive mechanisms provide parsimonious explanations for a number of puzzling phenomena and are compatible with proposed functions for CENP-B in enhancing chromosome segregation fidelity (Fachinetti, et al. 2015). They resolve the CENP-B paradox by suggesting a context-specific role for helix-deforming DNA-binding proteins in facilitating DNA secondary structure formation and/or transcription. In addition to recruiting HJURP to centromeres, non-B-form DNA and/or active transcription may suppress the unscheduled incorporation of canonical H3-containing nucleosomes (Nickol and Martin 1983) and explain the enrichment of DNA breaks and some damage repair proteins at centromeres (Guerrero, et al. 2010; Crosetto, et al. 2013) (Lu, et al. 2015). Consistent with the expansion of centromeres by HJURP tethering (Perpelescu, et al. 2015) and the high frequency of neocentromere formation in chicken (Shang, et al. 2013), we observed that the chicken genome is predicted to form non-B-form DNA structures more favorably than mammalian genomes.

This view also unifies the genetic and epigenetic conceptions of the centromere by explaining why CENP-B is necessary for *de novo* centromerization of artificial chromosomes (Ohzeki, et al. 2002), but not required to maintain native centromeres, which could be propagated by the presence of CENP-A (Fachinetti, et al. 2013) or Mis18 (Nardi, et al. 2016). Similarly, neocentromeres may be seeded in loci that adopt non-B conformations due to sequence and/or chromatin features. Centromere specification by recognition of nucleic acid structures would permit conservation of the general architecture of the centromere and kinetochore (Drinnenberg, et al. 2016) while providing a large sequence space that can be sampled during rapid evolution of DNA, suggesting a basis for driving genetic conflict at centromeres.

Therefore, genetically encoded structures may represent a common mechanism for specification of eukaryotic centromere identity.

## Author contributions

S.K. performed the analyses, S.H. performed the experiments, and S.K. and S.H. wrote the manuscript.

## Acknowledgements

We gratefully acknowledge P. Talbert, S. Ramachandran, J. Thakur, K Ahmad, and D. Melters for insightful discussions and suggestions and J. Henikoff for assistance with data analysis. We thank S. Biggins and H. Malik for comments on the manuscript. This work was funded by support from the Micki & Robert Flowers ARCS Endowment from the Seattle Chapter of the ARCS Foundation (S.K.) and the Howard Hughes Medical Institute (S.H.).

## Methods

<u>Datasets</u>. NCBI Sequence Read Archive accession numbers and references for publicly available datasets from a variety of species used in this study are included in **Table S1**. Note that members of the *Chlorocebus* and *Macaca* genera included in this study may represent subspecies rather than bona fide species (Yan, et al. 2011; Warren, et al. 2015). Illumina whole-genome sequencing (WGS) data selected were paired-end ~100x100-bp datasets to facilitate analysis of repeat variation.

<u>Pre-processing of Illumina data</u>. Raw paired-end Illumina reads were subjected to adapter trimming and quality filtering using BBDuk (http://jgi.doe.gov/data-and-tools/bbtools/) with the following parameters:

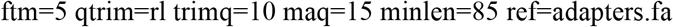

The FASTA file ‘adapters.fa’ is part of the BBDuk package and contains sequences of the Illumina TruSeq adapters. All subsequent analyses were performed on trimmed and filtered Illumina reads.

<u>Alignment of sequencing data</u>. Bowtie2 (v2.2.5) was used to perform alignments to published reference genomes or custom references as indicated. The following alignment parameters were used for paired-end reads:

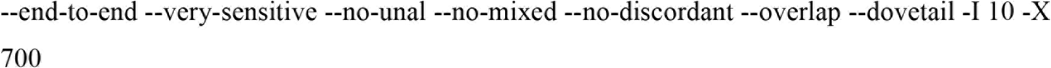

Alignment parameters used for single-end reads were:

--very-sensitive --no-unal --non-deterministic

<u>Reference genomes for short read alignment</u>. The following reference assemblies available from the UCSC Genome Browser were used: hg38 (human), mm10 (mouse), galGal5 (*Gallus gallus*), and sacCer2 (*S. cerevisiae*). For human and mouse, masked versions of the hg38 and mm10 assemblies were created using the hard-masked sequences available from the UCSC Genome Browser. For African green monkey, the RefSeq *Chlorocebus sabaeus* assembly (accession GCF_000409795.2) was hard-masked using RepeatMasker annotations available from RefSeq. The *S. pombe* assembly (ASM294v2) was downloaded from PomBase (McDowall, et al. 2015); the *S. mikatae* (IFO 1815^T^) and *S. kudriavzevii* (IFO 1815^T^) ultra-scaffolds were previously published (Scannell, et al. 2011) and are available online (http://sss.genetics.wisc.edu/cgi-bin/s3.cgi). The *S. castellii* assembly (NRRL-Y12630) was downloaded from the Saccharomyces Genome Database (Cherry, et al. 2012) and the *S. dairenensis* genome is available from the NCBI Assembly database (accession no. GCF_000227115.2). In all cases, Bowtie2 indexes were built using default parameters.

<u>*De novo* definition of centromeric satellite units</u>. Sanger reads, contigs from whole-genome assembly, and contigs from local assembly of Illumina reads were used to define centromeric satellites. Tandem Repeats Finder v5.02 (TRF) (Benson 1999) was used to identify all tandemly repeated sequences. Sequences corresponding to peaks in the resulting repeat length histograms that were not other abundant repeats (Alu, etc.) were classified as putative centromeric satellites. TRF was run with the following parameters: 2 7 7 80 10 50 1000 -h -ngs

Sequences from TRF peaks that passed a DUST complexity filter (implemented in PRINSEQ, http://prinseq.sourceforge.net; parameters: −lc_method dust −lc_threshold 7) were retained for subsequent analysis. In order to define unique monomers without shifting sequences to occupy similar registers, we took all tandem repeats corresponding to the major peak and subjected them to local alignment-based clustering using CD-HIT-EST (Li and Godzik 2006) with the following parameters: -c 0.8 -bak 1 -M 0 -d 0 -n 4 -G 0 -A 43

For each species, CD-HIT-EST-reported consensus sequences for clusters containing at least 1% of the input sequences were used to construct a BLAST database, which was then used to scan the Sanger reads and contigs and define new monomer locations. BLASTN searching was performed with the following options: -task blastn -num_alignments 1

<u>Identification of satellite monomer fragments in Illumina datasets.</u> Species-specific repeat databases produced as described above were used to identify fragments of monomers in paired-end Illumina sequencing datasets using BLASTN with the following options: -task blastn -num_alignments 1 -outfmt “6 qseqid qstart qend sseqid evalue sstrand pident length qlen”

The high depth of genome coverage in the selected datasets necessitated randomly sampling up to 10^6^ reads for each species.

<u>Principal components analysis (PCA) and clustering of Illumina reads</u>. We analyzed intra- and inter-specific variation in centromeric satellite sequences using dimensionality reduction. At least 10,000 satellite-containing reads were sampled from each species and oriented relative to the species-specific consensus sequences based on the BLASTN-reported alignment strand (see above). For each read, a *k*-mer frequency vector was created by sliding a *k*-length window in 1-nt steps along the length of the read and enumerating the fractional representation of each observed *k*-mer. For each species, *k*-mers appearing in more than 10 reads and in less than 50% of reads were retained for subsequent analysis. PCA was then performed on the set of frequency vectors for each species (**Figure S1C,D**). To account for the possibility of clusters with non-convex topology, 2000 reads used for PCA analysis were subjected to spectral clustering with number of clusters varying from 1 to 19. The silhouette coefficient (Rousseeuw 1987) was computed for each clustering. The clustering that produced the maximum silhouette coefficient was selected as the optimal clustering for each species and *k*-mer length (**Figure S1C,D**).

<u>CENP-B box abundance in primates and mouse.</u> To estimate the abundance of CENP-B boxes, occurrences of matches to the multiple alignment consensus CENP-B box sequence (TTCGNNNNANNCGGG), which is required for DNA-binding, were counted in Illumina WGS data. We further used the SELEX-defined CENP-B motif (Jolma, et al. 2013)(JASPAR accession MA0637.1) to search for matches using FIMO (Grant, et al. 2011) with default parameters.

<u>CUT&RUN profiling of CENP-A and CENP-B in human and African green monkey cell lines.</u> CUT&RUN was performed as described (Skene and Henikoff 2017b, a). Briefly, K562 or Cos-7 cells were gently washed twice in room temperature Wash buffer [20 mM HEPES pH 7.5, 150 mM NaCl, 0.5 mM spermidine and a Roche complete EDTA-free tablet (Sigma-Aldrich) per 50 ml], with 3 min centrifugation at 600 xg, mixed with activated Concanavalin A-coated magnetic beads (Bangs Laboratories) and rotated 5-10 min. Beads were captured by placing on a magnet stand, decanted and resuspended in Antibody buffer [2 mM EDTA and antibody at 1:100 in Dig-wash (Wash buffer supplemented with 0.05% digitonin (Calbiochem)]. Antibodies used were CENP-A (Abcam ab13939, mouse monoclonal), CENP-B (Abcam ab25734), Histone H3K27me3 (Cell Signaling Technologies 9733), and IgG (either Antibodies Online ABIN102961 guinea pig anti-rabbit, or GeneTex GTX105137, rabbit anti-human mitochondrial RNA polymerase). After overnight binding at 4 °C with rotation, beads were captured and washed once or twice in Dig-wash. For the CENP-A samples, beads were resuspended in secondary antibody (Abcam Ab46540, Rabbit anti-mouse) in Dig-wash and incubated 1 hr at 4 °C and washed once in Dig-wash. Beads were resuspended in Protein A-MNase (Batch #5 360 μg/ml) 1:500 in Dig-wash, incubated 1 hr at 4 °C, washed twice in Dig-wash and resuspended in 100 μL Dig-wash. Tubes were placed 0 °C, mixed with 2 μL 100 mM CaCl_2_, incubated at 0 °C for 30 min, and reactions were stopped by addition of 100 μL 340 mM NaCl, 20 mM EDTA pH8, 4 mM EGTA, 0.05% digitonin, 50 μg/ml glycogen and 200 pg mono-nucleosomal *S. cerevisiae* (spike-in) DNA. Samples were incubated 10 min at 37 °C and centrifuged 5 min 16,000 xg at 4 °C and the supernatant was treated with 25 μg/ml RNAse A (Thermo) 10 min at 37 °C. After phenol-chloroform-isoamyl alcohol and chloroform extraction, DNA was precipitated by addition of 2.5 volumes 100% ethanol, chilled on ice, centrifuged 10 min at 4 °C at 16,000 xg and the pellets were rinsed in 100% ethanol and air-dried. Pellets were resuspended in 1 mM Tris pH8 0.1 mM EDTA and used for standard Illumina library preparation. Paired-end PE25x25 sequencing was performed by the Fred Hutch Shared Genomics Resource. Data have been deposited in GEO (GSEXXXXX).

<u>Analysis of CUT&RUN data.</u> CENP-A and CENP-B CUT&RUN data were aligned to human alphoid BAC reference sequences described previously (Henikoff, et al. 2015) or to an α-satellite-containing *Chlorocebus aethiops* BAC sequence (GenBank accession AC239401.3). Read length histograms were generated by counting the Bowtie2r-reported aligned paired-end fragment lengths and autocorrelation of the resulting read length distributions was performed using Numpy. H3K27me3 CUT&RUN data were aligned to repeat-masked versions of the hg38 and *Chlorocebus sabaeus* assemblies as described above.

<u>Detection of perfect dyad symmetries in Illumina reads</u>. A suffix array-based approach was used to detect dyad symmetries. Each sequence of interest was concatenated with its reverse complement and the suffix and longest common prefix (LCP) arrays were constructed. Traversal of the suffix array, which indexes the suffixes of the sequence and its reverse complement in lexicographic order, provides an efficient way to identify repeated patterns. Specifically, in the case of a pattern with dyad symmetry, the sequence on the forward and reverse strands should occupy adjacent positions in the suffix array. Dyad symmetries of various lengths (3-7 bp) separated by “spacer” sequences (0-20 bp) were discovered using this approach.

<u>Detection of perfect and imperfect dyad symmetries in Illumina reads and genomic regions</u>. We used EMBOSS palindrome (Rice, et al. 2000) to detect dyad symmetries with mismatches in the palindromic region with the following parameters (varying the number of mismatches):

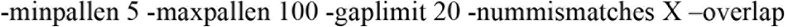

For each position in a sequence of interest, we defined dyad density as the sum of the lengths of the palindromic regions that contain that position. For a sequence, the length-normalized dyad density was defined as the sum of the per-position values divided by the sequence length.

<u>DNA secondary structure prediction</u>. RNAfold from the ViennaRNA package (Lorenz, et al. 2011) was used to predict folding free energies. RNAfold was used with the following parameters for DNA secondary structure prediction:

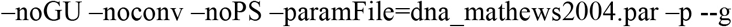

Predictions were performed on random samples of 10^4^ BLAST-defined monomers (Sanger data) or 10^4^ reads containing BLAST matches (Illumina data).

<u>Prediction of DNA melting and cruciform transitions</u>. Strand separation and cruciform extrusion propensities were predicted using SIST (Zhabinskaya, et al. 2015) with default parameters. For sequences greater than 10 kb in length (the maximum permissible length compatible with SIST), we slid a 5 kb window in 2.5 kb steps to generate short sequences that were analyzed using SIST. These SIST predictions were then reassembled for the full sequence by conservatively taking the maximum at each base (for bases spanned by multiple windows). For a given sequence, melt and cruciform scores were computed by summing the estimated transition probabilities for each position in the sequence and dividing by the length of the sequence.

<u>Selection of control genomic regions</u>. To account for sequence composition of centromere and neocentromere sequences, we selected random genomic regions without known centromere activity and with similar dinucleotide composition. Dinucleotide frequencies for a query sequence of interest were calculated and the Spearman rank correlation (ρ) was used to identify non-overlapping windows of the same length as the query in a genome with a similar dinucleotide frequency pattern. We considered two regions to be sufficiently similar if ρ ≥ 0.9 and excluded regions overlapping annotated centromeric sequences. Up to 1,000 random sites defined using this procedure were used for comparisons.

<u>Statistical analyses.</u> The two-sample Kolmogorov-Smirnov test was used to compare distributions of values of interest *(e.g.,* dyad density, SIST scores).

<u>Analysis of CENP-A ChIP-seq data</u>. Our published CENP-A ChIP-seq 100x100 bp Illumina data were subjected to the same adapter trimming and quality filtering steps described above prior to merging pairs using SeqPrep as described previously (Henikoff, et al. 2015). Merged pairs were aligned to alphoid reference sequences using Bowtie2 using the single-end mapping parameters described above. Data from ChIP-seq of *S. cerevisiae* Cse4 (Henikoff, et al. 2014), *S. pombe* Cnp1 (Thakur, et al. 2015), and CENP-A in human neocentromeres cell lines (Hasson, et al. 2013) were aligned using the paired-end mapping parameters described above.

<u>Analysis of DNase-seq data</u>. Adapter trimmed and quality filtered human and mouse paired-end DNase-seq (DNase-FLASH) ENCODE datasets were aligned to Sanger reads containing satellite sequences. Fragment length vs. midpoint plots (V-plots) were constructed as previously described (Henikoff, et al. 2011).

<u>Analysis of ssDNA-seq data</u>. Raw reads were quality filtered and subjected to adapter trimming as described above.

<u>Analysis of motif similarity</u>. Tomtom v4.12.0 (Gupta, et al. 2007) with default parameters was used to determine similarity of yeast CDEI-derived motifs to a library of consensus sites for 203 yeast transcription factors (MacIsaac, et al. 2006) and to a Reb1 motif defined using high-resolution ChIP-seq (Kasinathan, et al. 2014).

<u>Source code</u>. Code for performing the described analyses is available from Github (https://github.com/sivakasinathan/cenpb).

## Supplementary Figures & Tables

**Figure S1.**
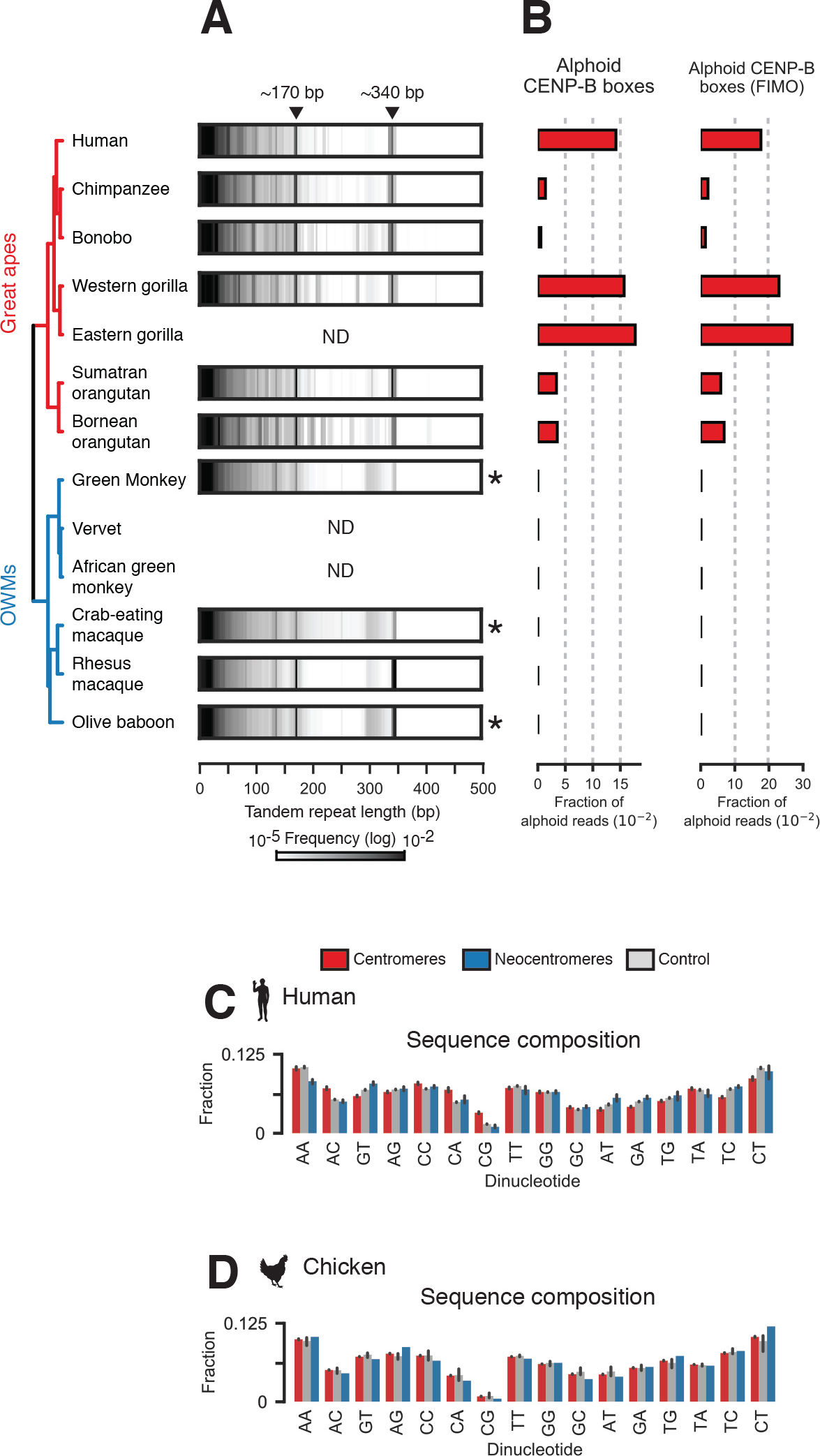
Characterization of variation in vertebrate centromeres, Related to Figures 1,2,4, and 5. **(A)** Heatmap representations of histograms of lengths of tandem repeats discovered *de novo* in datasets for great apes and Old World Monkeys (OWMs). Sequences from indicated peaks at ~170-bp and ~340-bp were used to generate species-specific repeat libraries. Where available, raw shotgun Sanger reads were used. If Sanger reads were unavailable, the genome assembly (including unplaced contigs) was scanned for tandem repeats (indicated by an asterisk); “ND” indicates neither Sanger reads nor an assembly were available. (**B**) Fraction of alphoid reads containing matches to the SELEX-defined CENP-B box motif deposited in JASPAR. Examples of dinucleotide profiles for centromeric regions, neocentromeres, and non-centromeric background genomic regions selected by *k*-mer matching from human (**C**) and chicken (**D**).

**Figure S2.**
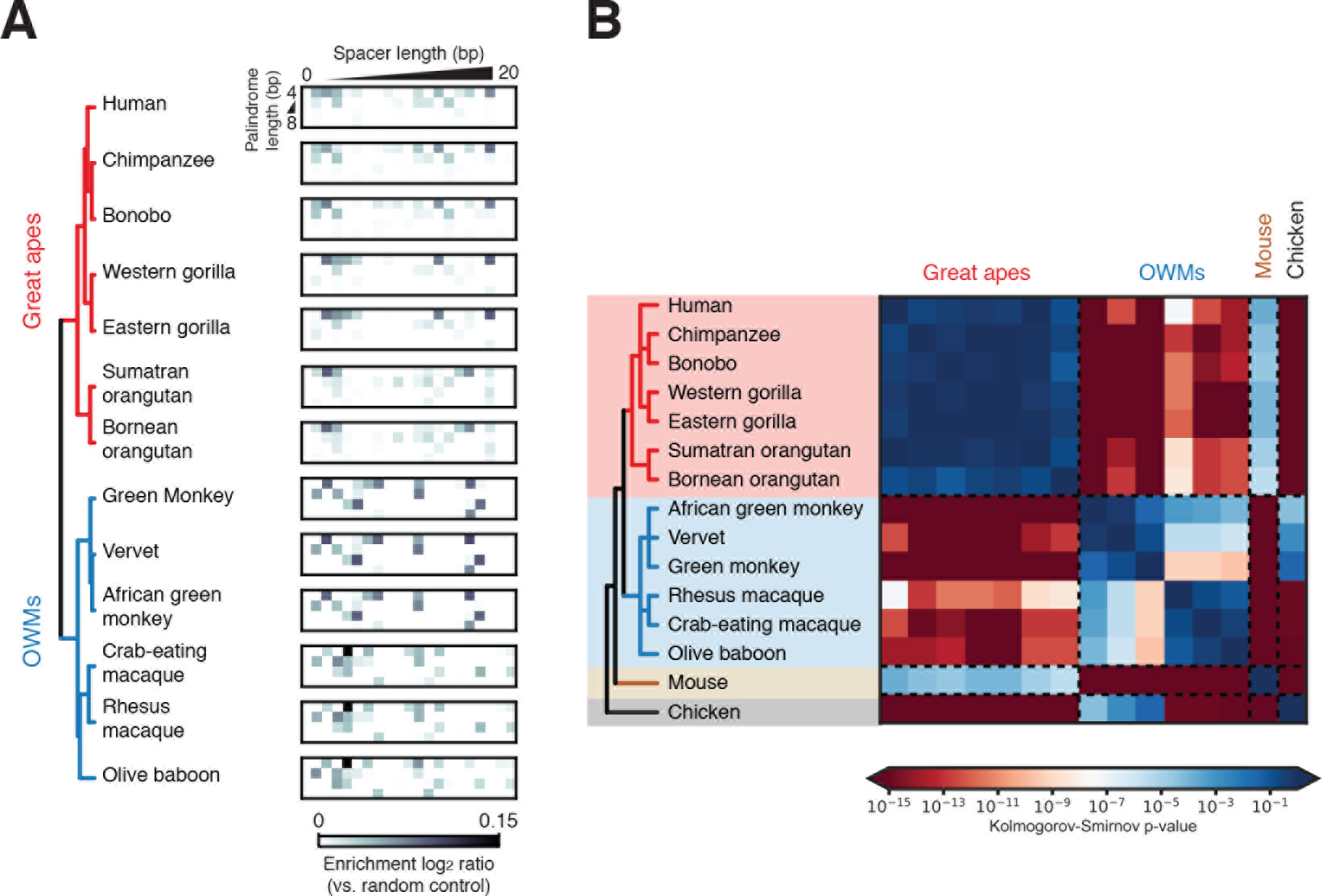
Clade-specific secondary structure-forming dyad symmetries at vertebrate centromeres, Related to Figures 1 and 2. (**A**) Enrichment (vs. permuted sequence) of dyad symmetries with varying palindrome and spacer lengths for great apes and Old World Monkeys (OWMs) allowing one mismatch in the palindromic region. (**B**) Two-sample Kolmogorov-Smirnov *p*-values for all pairwise comparisons of distributions of RNAfold-predicted ensemble free energies for great ape, OWM, mouse, and chicken centromeric sequences. Heatmap depicts -log(*p*-value) with warmer (red) colors indicating dissimilarity and colder (blue) colors indicating similarity.

**Figure S3.**
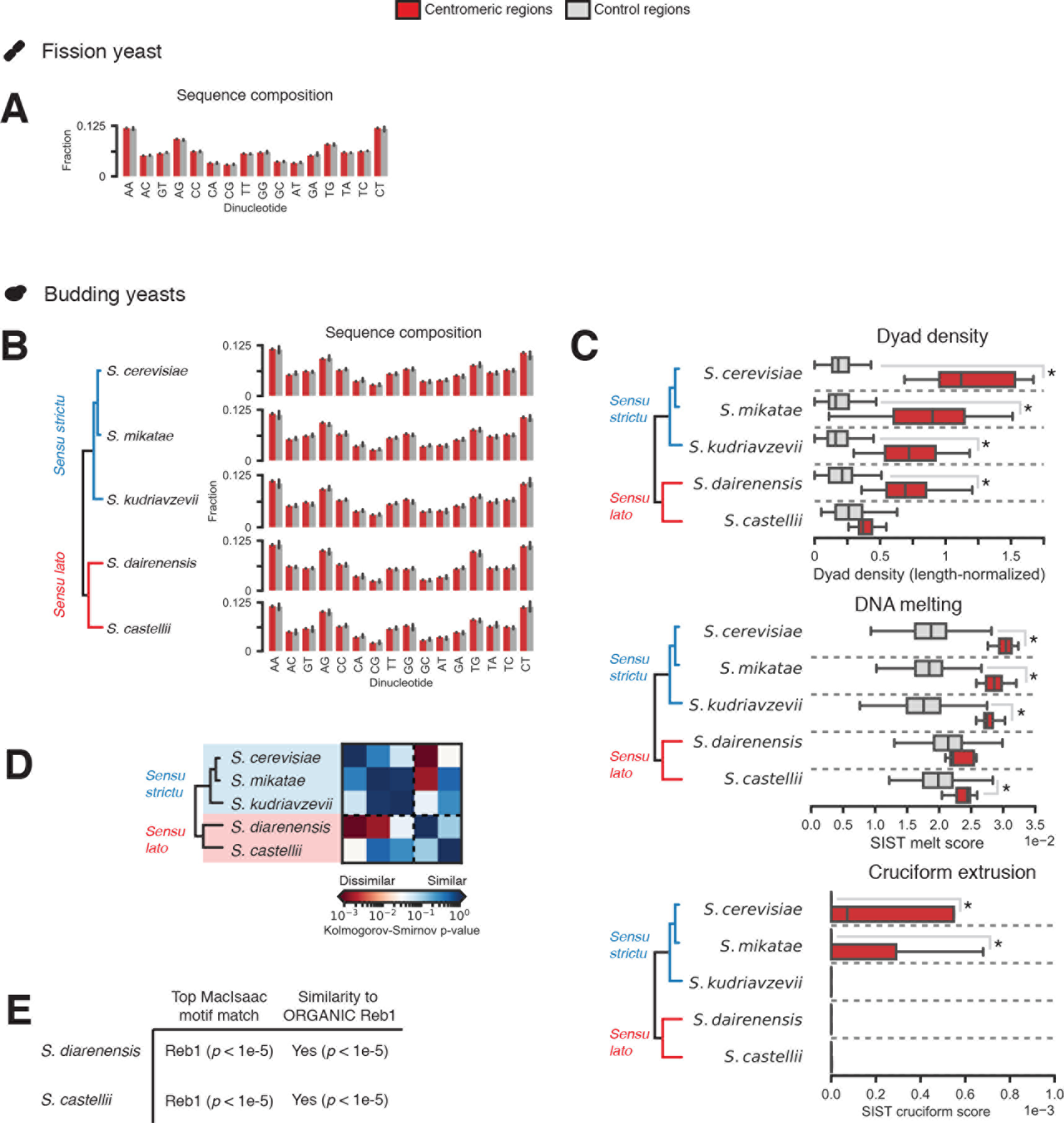
Analysis of yeast centromeric DNAs, Related to Figure 6. (**A**) Dinucleotide composition at *S. pombe* centromeres and matched non-centromeric background sregions sampled from the fission yeast genome assembly. (**B**) Dinucleotide composition at centromeres and randomly-selected, matched non-centromeric regions from the genomes of selected *sensu strictu* and *sensu lato* saccharomycete genomes. (**C**) Dyad symmetry density and DNA melting and cruciform extrusion predilections at centromeres and composition-matched non-centromeric random regions from the genomes of selected saccharomycetes. Asterisks indicate two-sample Kolmogorov-Smirnov *p* < 0.05. (**D**) Two-sample Kolmogorov-Smirnov *p*- values for all pairwise comparisons of RNAfold-predicted ensemble free energies for centromeric sequences from saccharomycetes. (**E**) Tomtom motif similarity analysis of CDEI motifs from two *sensu lato* yeasts with the highest scoring hit from scanning a database of binding sites for ~200 yeast transcription factors (MacIsaac, et al. 2006) and comparison to the Reb1 binding site defined by ORGANIC profiling, a high-resolution mapping approach (Kasinathan, et al. 2014).

**Table S1.**
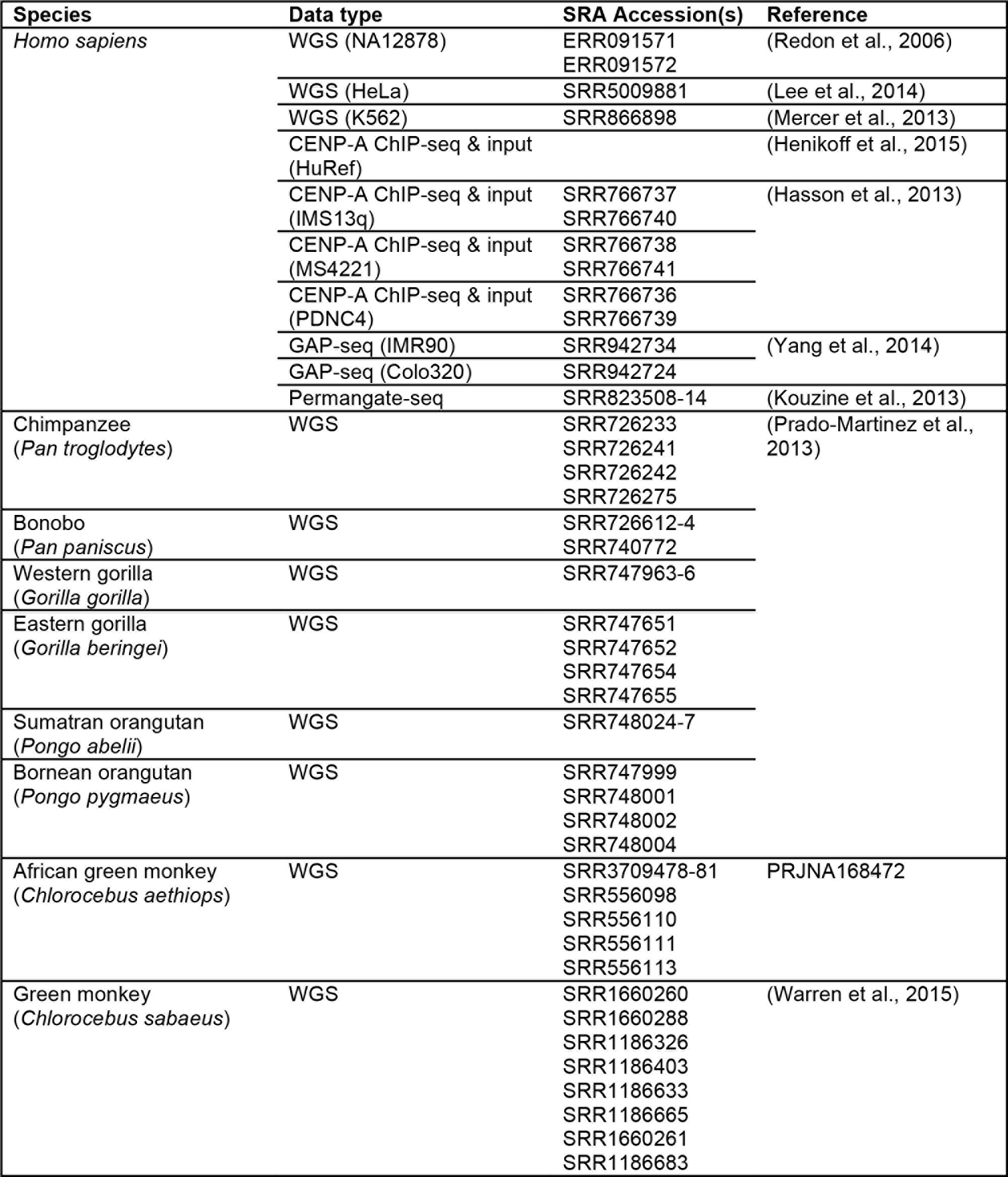
Publicly available datasets analyzed in this study, Related to STAR Methods. NCBI Sequence Read Archive (SRA) accession numbers and references for whole-genome (WGS), chromatin immunoprecipitation (ChIP-seq), genome-wide analysis of palindrome formation (GAP-seq), permanganate-seq, and DNase-seq Illumina datasets are provided. Where references are unavailable, NCBI BioProject accessions for public reference genome projects are provided.

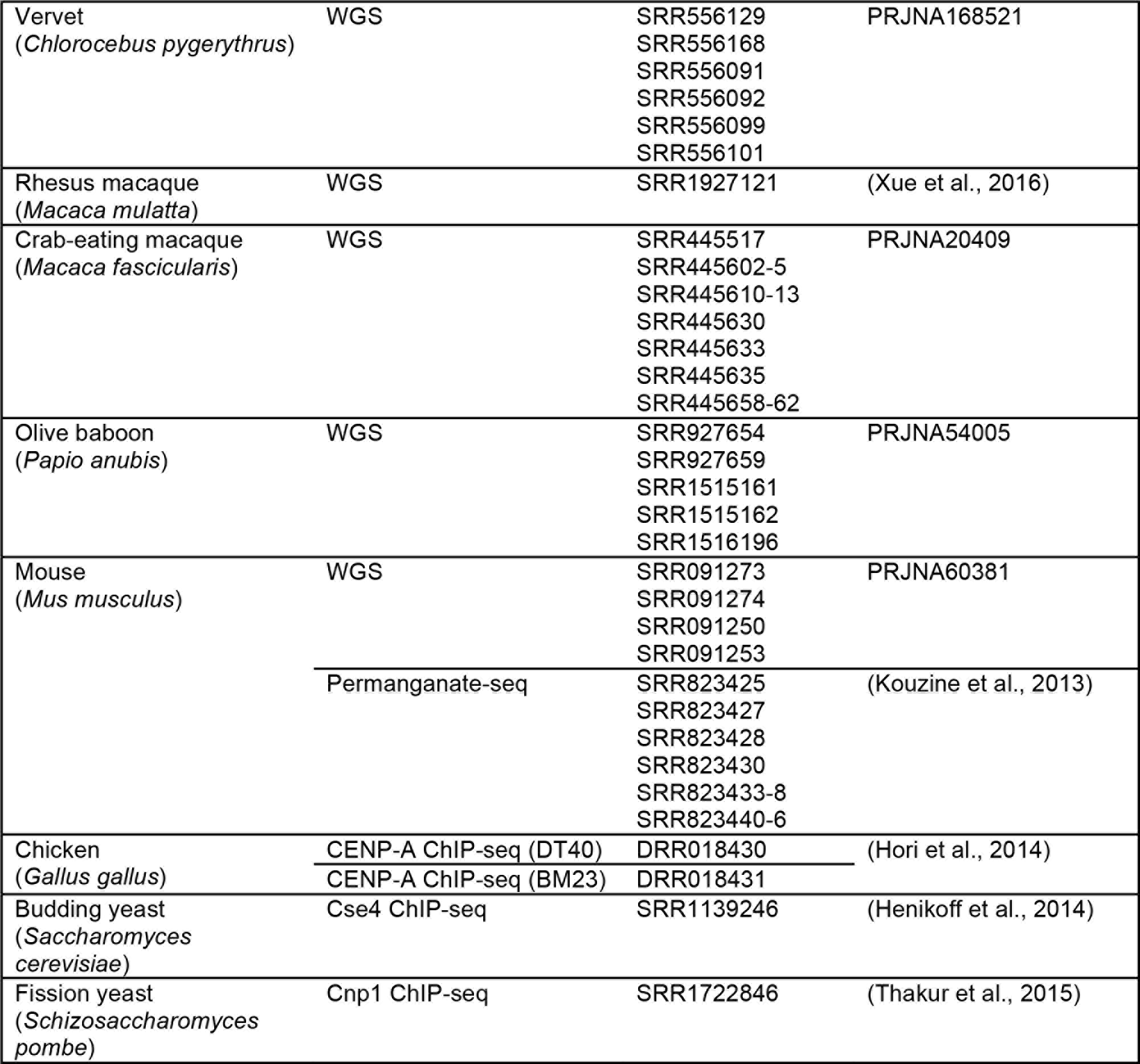

